# Oleuropein stimulates peripheral serotonin secretion via voltage-dependent Ca^2+^ channels and acutely regulates central function

**DOI:** 10.64898/2026.07.15.738825

**Authors:** Yuki Kamei, Rina Sugitani, Noa Onzawa, Mana Akao, Taiki Fushimi, Mitsugu Akagawa

## Abstract

Serotonin (5-HT) is a monoamine which regulates not only central functions but also various peripheral functions. Peripheral 5-HT is primarily derived from the gut, and its synthesis and secretion are regulated by enterochromaffin cells. Stimulation of enterochromaffin cells by food factors may regulate mental functions via the gut-brain axis. We studied whether oleuropein, an olive-derived polyphenol, regulates central functions by stimulating 5-HT secretion from enterochromaffin cells. In QGP-1 cells, which are enterochromaffin-like cells, 10 µM oleuropein stimulated 5-HT secretion via Ca^2+^ influx through T-type and L-type voltage-dependent Ca^2+^ channels. Hydroxytyrosol, a metabolite of oleuropein, also promoted 5-HT secretion via the same mechanism. Furthermore, oleuropein stimulated 5-HT secretion from isolated mouse colon via voltage-dependent Ca^2+^ channels. Finally, oral administration of 200 mg/kg oleuropein acutely increased the depression-like behavior in the mice, which was inhibited by the prior administration of ramosetron, a 5-HT_3_ receptor antagonist. These findings suggest that oleuropein is a potent stimulant of gut 5-HT secretion, and show that food factors may act to regulate mental function via the secretion of gut 5-HT.

## 1. INTRODUCTION

Serotonin (5-hydroxytryptamine or 5-HT) a monoamine synthesized from L-tryptophan via hydroxylation by tryptophan hydroxylase (TPH) and decarboxylation by aromatic L-aromatic amino acid decarboxylase, regulates diverse biological functions.^1^ The rate-limiting enzyme in 5-HT synthesis is TPH which exists in two subtypes: TPH1 is abundant in the gut, and TPH2 is abundant in the brain.^2^ Gut-derived 5-HT regulates gastrointestinal motility and is also taken up by platelets, circulating systemically to control peripheral functions such as vascular functions and metabolism via various 5-HT receptors.^3,4^ In contrast, brain-derived 5-HT regulates diverse central functions, such as behavior, appetite, and thermoregulation, through serotonergic signaling and modulation of gamma amino butyric acidergic, dopaminergic, and noradrenergic neural circuits.^5^ Although gut-derived 5-HT cannot cross the blood-brain barrier,^6^ recent studies have suggested that gut-derived 5-HT may also regulates these central functions. For example, TPH1 knockout rats exhibited decreased anxiety-like behavior even though only peripheral 5-HT levels were reduced.^7^ Furthermore, selective activation of enterochromaffin cells, which are responsible for the synthesis and secretion of 5-HT in mice, has been suggested to leads to the exhibition of anxiety-like behavior.^8^ These reports suggest that gut-derived 5-HT may serve as a potential therapeutic target not only for various peripheral diseases but also for mental disorders as a key driver of the gut-brain axis.

Enterochromaffin cells are a type of neuroendocrine cell sparsely distributed within the intestinal epithelium and play a crucial role in regulating 5-HT function.^9^ Enterochromaffin cells sense various stimulants in the intestinal lumen, such as food factors, gut bacteria-derived metabolites, and enterotoxins. These stimulants promote Ca^2+^ influx via many types of Ca^2+^ channels,^10,11^ and an increase in cytosolic Ca^2+^ level stimulates the release of 5-HT stored in vesicles via typical Ca^2+^-dependent exocytosis.^12,13^ Among the factors that stimulate 5-HT secretion, food factors are easy to include in the daily diet and may contribute to the regulation of central functions mediated by gut 5-HT. Many food factors act as ligands for transient receptor potential (TRP) channels, the sensory receptors, and increase cytosolic Ca^2+^ levels.^14^ For example, cinnamaldehyde is known to stimulate 5-HT secretion via TRPA1.^15^ However, very few food factors have been proven to stimulate gut 5-HT secretion and exert peripheral or central effects.

Oleuropein, a polyphenol found in olive fruit and leaves, is composed of tyrosol, elenolic acid, and glucose (Figure 1A).^16^ Both oleuropein and its metabolite, hydroxytyrosol, have potent antioxidant effects and may contribute to various health benefits associated with olive consumption.^17,18^ Olive consumption is associated with improvements of pathological conditions involving 5-HT, such as functional gastrointestinal disorders and mental disorders,^19,20^ but the role of oleuropein in these effects has not been fully investigated. Interestingly, oleuropein has been reported to increase cytosolic Ca^2+^ concentrations in some cell types, suggesting that it may be a stimulant for 5-HT secretion.^21^ ^22^ However, no studies have investigated the effect of oleuropein on 5-HT secretion from enterochromaffin cells. In the present study, we aimed to investigate whether oleuropein regulates central functions by stimulating 5-HT secretion from enterochromaffin cells.

**Figure 1.**
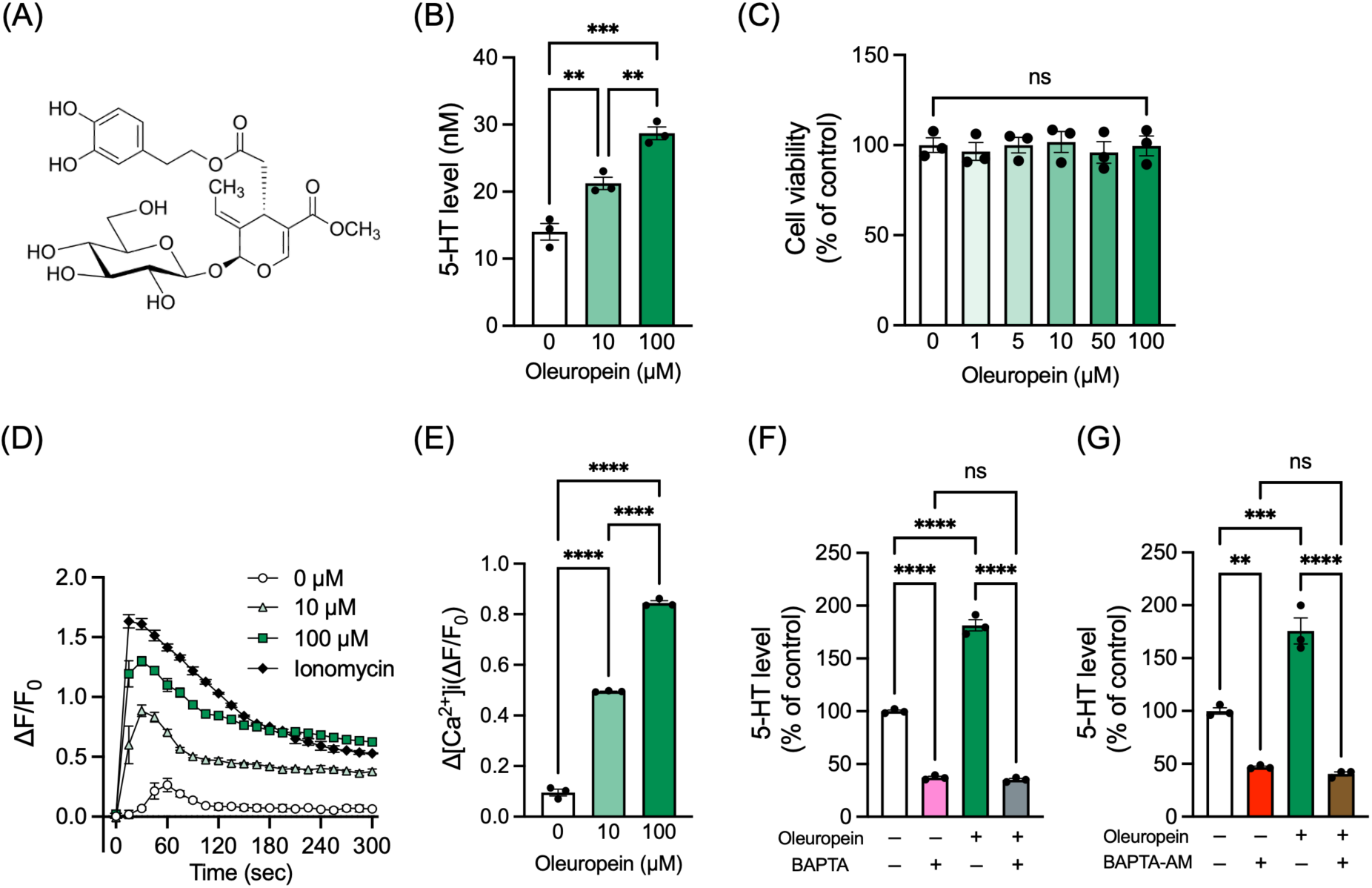
Effects of oleuropein on 5-HT secretion from enterochromaffin-like cells and Ca^2+^ influx. (**A**) Structure of oleuropein. (**B**) Dose-dependent effects of oleuropein on 5-HT secretion. QGP-1 cells were incubated with HBSS containing different concentrations of oleuropein and the supernatant was subjected to HPLC analysis. (**C**) Dose-dependent effects of oleuropein on cell viability. QGP-1 cells were incubated with HBSS containing different concentrations of oleuropein for 60 min and subjected to the WST-8 assay. (**D**) Dose-dependent effects of oleuropein on cytosolic Ca^2+^ levels. Fluo-8 AM was internalized into QGP-1 cells, and oleuropein was added. The fluorescence intensity was then measured over time for 5 min. Ionomycin was used as a positive control. (**E**) Quantification of changes in the Fluo-8 fluorescence intensity. The average change in fluorescence intensity from 0 min after oleuropein addition to each time point was expressed as Δ[Ca^2+^]i (ΔF/F0). (**F**) Effects of cell-membrane-impermeable Ca^2+^ chelator on 5-HT secretion by oleuropein. QGP-1 cells were incubated with HBSS containing 100 µM oleuropein for 5 min with or without BAPTA, and the supernatant was subjected to HPLC analysis. (**G**) Effects of cell-membrane-permeable Ca^2+^ chelator on 5-HT secretion by oleuropein. QGP-1 cells were incubated with HBSS containing 100 µM oleuropein for 5 min with or without BAPTA-AM and the supernatant was subjected to HPLC analysis. Bars represent the mean ± SEM, and dots indicate individual data points (*n* = 3). \*\**p* < 0.01, \*\*\**p* < 0.001, and \*\*\*\**p* < 0.0001, one-way ANOVA, Tukey’s multiple comparisons test. ns denotes not significant.

## 2. MATERIALS AND METHODS

### 2.1 Cell culture

Human somatostatinoma-derived QGP-1 cells were obtained from JCRB Cell Bank (JCRB0183) and used as enterochromaffin-like cells. Cells were cultured in RPMI 1640 (Nacalai Tesque) with 5% fetal bovine serum (FBS, Sigma-Aldrich), 100 U/mL penicillin, and 100 µg/mL streptomycin (FUJIFILM Wako Pure Chemical) at 37 ℃ in an atmosphere containing 5% CO_2_.

### 2.2 Cell treatment and sample collection

After confirming subconfluence, cells were washed with Hank’s balanced salt solution containing Ca^2+^ (HBSS (+), FUJIFILM Wako Pure Chemical) twice and treated with HBSS (+) containing 1-100 µM oleuropein (Tokyo Chemical Industry) or hydroxytyrosol (BLD Pharmatech) or 0.1% dimethyl sulfoxide (DMSO) as vehicle for 5 min. For studying the involvement of Ca^2+^ in 5-HT secretion, 600 µM 1,2-Bis(2-aminophenoxy)ethane-*N,N,N’,N*’-tetraacetic acid (BAPTA, Tokyo Chemical Industry), an extracellular Ca^2+^ chelator, or 25 µM BAPTA-acetoxymethyl ester (BAPTA-AM, Invitrogen), a cell membrane-permeable Ca^2+^ chelator, or 0.1% DMSO was additionally added to the corresponding group. BAPTA was treated immediately before the addition of the food factor, while BAPTA-AM was treated for 30 min prior to the addition of the food factor. For identification of Ca^2+^ channels involved in 5-HT secretion, 30 µM HC-030031, a TRPA1 inhibitor (FUJIFILM Wako Pure Chemical), 30 µM capsazepine, a TRPV1 inhibitor (Nacalai Tesque), 5 µM NNC 55-0396, a T-type voltage-dependent Ca^2+^ channel inhibitor (TargetMol), or 100 µM nifedipine, a L-type voltage-dependent Ca^2+^ channel inhibitor (FUJIFILM Wako Pure Chemical) was additionally added to the corresponding group. The culture supernatant was collected and centrifuged at 1,000 ×g, 4 ℃ for 5 min to remove cell debris. The supernatant was collected and stored at -20 ℃ until high performance liquid chromatography (HPLC) analysis.

### 2.3 Measurement of 5-HT by HPLC

The culture supernatant was analyzed by HPLC using an HPLC pump (L-2130, HITACHI) equipped with an HPLC column (COSMOSIL 5C_18_-AR-Ⅱ, 4.6 mm I.D.×250 mm, Nacalai Tesque) and fluorescence detector (L-7485, HITACHI, Ex/Em = 492/530 nm). The mobile phase consisted of 50 mM KH_2_PO_4_ (pH 5.0)/acetonitrile (95:5, *v/v*), and the flow rate was 1.0 mL/min. The sample injection volume was 20 µL. 5-HT standard was prepared using serotonin hydrochloride (Nacalai Tesque). The 5-HT levels obtained from the supernatant immediately after the addition of the food factors were subtracted as a blank.

### 2.4 Evaluation of cytotoxicity

The cell viability was determined using the water soluble tetrazolium-8 (WST-8) assay to determine the cytotoxicity of oleuropein and hydroxytyrosol,. Cells were seeded into a 96-well plate and after confirming subconfluence, the culture supernatant was removed, HBSS was added, and oleuropein or hydroxytyrosol was then added to achieve a final concentration of 1–100 µM. After incubation for 1 h at 37 ℃ in an atmosphere containing 5% CO_2_, the culture supernatant was removed and 5% Cell Count Reagent SF (Nacalai Tesque) in RPMI 1640 without FBS was added to each well. After incubation for 2 h at 37 ℃ in an atmosphere containing 5% CO_2_, the plate was placed in a microplate reader (SH-1000, Corona), and the absorbance was measured at 450 nm.

### 2.5 Evaluation of cytosolic Ca^2+^ levels

The cell membrane-permeable fluorescent Ca^2+^ indicator Fluo-8 AM (AAT Bioquest) was utilized to evaluate the cytosolic Ca^2+^ levels. Cells were seeded in a 96-well plate which was coated with Cellmatrix TypeⅠ-C (Nitta Gelatin). After subconfluence was confirmed, the cells were washed with the assay buffer (NaCl: 137 mM, KCl: 5.4 mM, MgCl_2_: 2.5 mM, CaCl_2_: 2.3 mM, glucose: 100 mM, HEPES: 10 mM) and incubated for 40 min with assay buffer containing 3 µM Fluo-8 AM at 37 ℃ in an atmosphere containing 5% CO_2_. After Fluo-8 AM was incorporated into the cells, the plate was placed in a microplate reader (Synergy H1, Agilent) at 37 °C for 20 min to measure the baseline fluorescence intensity. Then, oleuropein, hydroxytyrosol or ionomycin (Alomone Labs) was added as the positive control, and the fluorescence intensity (Ex/Em = 492/530 nm) was measured every 15 sec for 300 sec. The involvement of voltage-dependent Ca^2+^ channels was investigated by treatment of 5 µM NNC 55-0396 or 100 µM nifedipine 5 min before adding food factors to the corresponding group. The average change in the fluorescence intensity from 0 min after food factors addition to each time point was expressed as Δ[Ca^2+^]i (ΔF/F0) and compared between groups.

### 2.6 Immunofluorescence staining

A glass slide with a paraffin section of the Swiss-rolled digestive tract from 8-week-old male C57BL/6 mouse was obtained from GenoStaff. The section was deparaffinized in xylene (Nacalai Tesque), rehydrated through a graded ethanol series, and antigen-activated by heating in a microwave oven with citrate buffer (10 mM citrate, pH 6.0). The section was rinsed with water and treated with Blocking One (Nacalai Tesque) to prevent nonspecific binding and then washed with TBS containing 0.05% Tween 20 (TBS-t). The section was incubated at 4 ℃ overnight with the primary antibody for chromogranin A (Proteintech, 1/1000) and for 5-HT (Imuunostar, 1/1000) diluted in Can Get Signal Solution 1 (Toyobo). The section washed with TBS-t was incubated at room temperature for 30 min with secondary antibody for goat anti-mouse conjugated with Cy2 (Chemicon, 1/200), and goat anti-rabbit conjugated with Cy3 (Amersham Biosciences, 1/1000) diluted in Can Get Signal Solution 2 (Toyobo). The section washed with TBS-t, was covered with a mounting medium containing DAPI (Nacalai Tesque). Fluorescence images were acquired using the EVOS™ M5000 imaging system equipped with LED Light Cubes for DAPI, GFP, and RFP and a 40x objective lens (Thermo Fisher Scientific).

### 2.7 Animal experiment

Five-week-old male C57BL/6J mice (Japan SLC) were housed under specific pathogen-free conditions and maintained under a 12 h light-dark cycle with free access to water and standard chow (MF diet, Oriental Yeast). The present study was approved by the Animal Experimentation Committee of Tokushima University School of Medicine (animal ethical clearance No. T2022–93) and was carried out in accordance with guidelines for the Animal Care and Use Committee of Tokushima University Graduate School.

### 2.8 *Ex vivo* culture of colon tissue

After acclimation for a week, the mice were euthanized by cervical dislocation, and the colon tissues were dissected and washed with phosphate buffer saline to remove the feces from the lumen. The proximal side of the tissue (30-50 mg) was isolated and washed by gently swirling in HBSS containing 2 µM fluoxetine, a selective 5-HT reuptake inhibitor. The colon tissues were washed and then incubated for 15 min with HBSS containing 2 µM fluoxetine with or without 100 µM oleuropein. For investigation of the involvement of voltage-dependent Ca^2+^channels, 5 µM NNC 55-0396 or 100 µM nifedipine were co-treated in the corresponding group. The culture supernatant was centrifuged and then subjected to HPLC analysis in the same manner as the cell culture supernatant described above.

### 2.9 Behavioral tests

After acclimation for a week, the mice were further acclimated in a quiet testing room maintained at constant conditions of 23-24 ℃ and 30-40% humidity for over 1 h before starting the behavioral tests. Ramosetron (Selleck), a 5-HT_3_ receptor inhibitor, or water as vehicle were administered orally to the respective groups 1 h prior to the start of the behavioral tests, and 200 mg/kg oleuropein or water as vehicle were administered orally to the respective groups 30 min prior to the start of the behavioral tests. The behavioral tests were conducted consecutively in order of least stress: open field test, tail suspension test, and forced swim test, with 10 min intervals between each test.

#### 2.9.1 Open field test

A single mouse was placed in the center of an open, walled arena (45 cm square, a matte white chamber made of vinyl chloride), and its locomotor activity was recorded from above for 5 min. The video was analyzed using Mouse Activity^23^, a MATLAB-based open-source video tracking program to calculate the total distance traveled and the time spent in the outer area of the arena. After each test, the arena was wiped down with 70% ethanol to eliminate any residual odors.

#### 2.9.2 Tail suspension test

The mice were suspended by attaching tape to the tip of their tails to ensure a height of 40 cm above the ground. To prevent the mouse from grabbing its own tail and affecting the activity time, a 4 cm long hose with an inner diameter of 4 mm was placed at the base of the tail. The activity was recorded for 6 min, and the immobility time during the 5 min period excluding the first 1 minute was accumulated.

#### 2.9.3 Forced swim test

A single mouse was placed into an acrylic pipe with a base measuring 10 cm in diameter and 25 cm in height, filled with water at 24–25 °C to a height of 16.5 cm. The activity was recorded for 6 min, and the immobility time during the 5 min period excluding the first 1 min was accumulated. Kicking the wall to maintain the body position was considered as immobility.

### 2.10 Statistical analysis

Statistical analysis was performed using one-way ANOVA with Tukey’s multiple comparisons test. All data analysis was performed using GraphPad Prism 11 software (Graphpad Software). *P* < 0.05 was considered statistically significant.

## 3. RESULTS

### 3.1 Oleuropein stimulates 5-HT secretion via Ca^2+^ influx

First, we investigated whether oleuropein enhances 5-HT secretion using pancreatic-derived QGP-1 cells, which are known as enterochromaffin-like cells.^24^ We confirmed that oleuropein promoted 5-HT secretion in a dose-dependent manner at 10 and 100 µM (Figure 1B) and that it was not due to 5-HT leakage through cytotoxicity (Figure 1C). Since 5-HT secretion is promoted by an increase in cytosolic Ca^2+^, we evaluated the immediate changes in cytosolic Ca^2+^ levels after oleuropein addition using Fluo-8, a fluorescent Ca^2+^ indicator, and found that the fluorescence intensity increased immediately after oleuropein addition (Figure 1D, E). To confirm whether Ca^2+^ influx is involved in the 5-HT secretion-promoting effect of oleuropein, we used cell-membrane-impermeable or cell membrane-permeable Ca²⁺ chelators. The results showed that the 5-HT secretion-promoting effect of oleuropein was completely abolished regardless of the Ca²⁺ chelator used (Figure 1F, G). These results indicate that oleuropein stimulates 5-HT secretion via Ca²⁺ influx.

### 3.2 5-HT secretion by oleuropein is mediated by voltage-dependent Ca²⁺ channels, but not TRP channels

Next, we attempted to identify the Ca²⁺ channels that mediate oleuropein-induced 5-HT secretion. First, we assumed the involvement of TRPA1, which is expressed in enterochromaffin cells and is affected by many food factors.^10,25^ However, the use of HC-030031, an inhibitor of TRPA1, did not diminish the 5-HT-secreting effect of oleuropein (Figure 2A). In addition, the use of capsazepine, an inhibitor of TRPV1, did not diminish the 5-HT-secreting effect of oleuropein (Figure 2B). Next, because oleuropein was reported to act on T-type voltage-dependent Ca²⁺ channel,^21^ we used NNC 55-0396, an inhibitor of the T-type voltage-dependent Ca²⁺ channel. The result showed that the inhibition of T-type voltage-dependent Ca²⁺ channel significantly diminished the 5-HT-secreting effect of oleuropein as well as the rise in cytosolic Ca²⁺ (Figure 2C, D). Furthermore, we used nifedipine, an inhibitor of L-type voltage-dependent Ca²⁺ channel since L-type voltage-dependent Ca²⁺ channel has been reported to be expressed on the apical side of enterochromaffin cells.^26,27^ Similar to the T-type voltage-dependent Ca²⁺ channel, inhibition of L-type voltage-dependent Ca²⁺ channel also diminished the 5-HT-secreting effect of oleuropein as well as the rise in cytosolic Ca²⁺ (Figure 2E to F). These results indicate that the 5-HT-secreting effect of oleuropein is mediated by Ca²⁺ influx via voltage-dependent Ca²⁺ channels, but not TRP channels.

**Figure 2.**
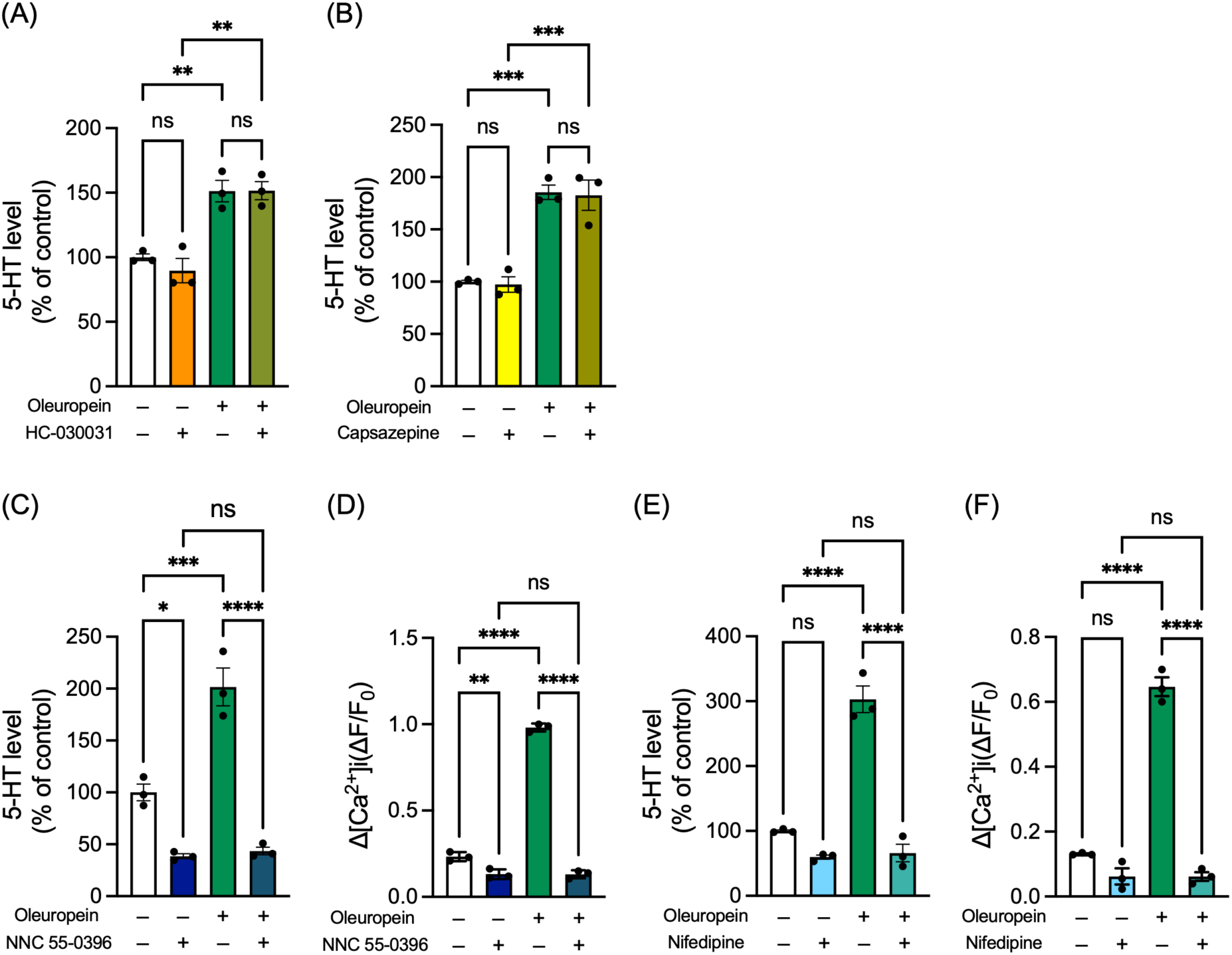
Effects of TRP channel inhibitors and voltage-dependent Ca^2+^ channel inhibitors on 5-HT secretion and Ca^2+^ influx induced by oleuropein. (**A**) Effect of TRPA1 inhibitor on 5-HT secretion by oleuropein. QGP-1 cells were incubated with HBSS containing 100 µM oleuropein for 5 min, with or without HC-030031. (**B**) Effects of TRPV1 inhibitor on 5-HT secretion by oleuropein. QGP-1 cells were incubated with HBSS containing 100 µM oleuropein for 5 min, with or without capsazepine. (**C**) Effects of T-type voltage-dependent Ca^2+^ channel inhibitor on 5-HT secretion by oleuropein. QGP-1 cells were incubated with HBSS containing 100 µM oleuropein for 5 min, with or without NNC 55-0396. (**D**) Effects of T-type voltage-dependent Ca^2+^ channel inhibitor on Ca^2+^ influx induced by oleuropein. Fluo-8 AM was internalized into QGP-1 cells, and oleuropein was added with or without NNC 55-0396. The fluorescence intensity was then measured over time for 5 min, and the average change in fluorescence intensity from 0 min after oleuropein and NNC 55-0396 addition to each time point was expressed as Δ[Ca^2+^]i (ΔF/F0). (**E**) Effects of L-type voltage-dependent Ca^2+^ channel inhibitor on 5-HT secretion by oleuropein. QGP-1 cells were incubated with HBSS containing 100 µM oleuropein for 5 min, with or without nifedipine. (**F**) Effects of T-type voltage-dependent Ca^2+^ channel inhibitor on Ca^2+^ influx induced by oleuropein. Fluo-8 AM was internalized into QGP-1 cells, and oleuropein was added with or without nifedipine. The fluorescence intensity was then measured over time for 5 min, and the average change in fluorescence intensity from 0 min after oleuropein and nifedipine addition to each time point was expressed as Δ[Ca^2+^]i (ΔF/F0). Bars represent the mean ± SEM, and dots indicate individual data points (*n* = 3). \**p* < 0.05, \*\**p* < 0.01, \*\*\**p* < 0.001, \*\*\*\**p* < 0.0001, one-way ANOVA, Tukey’s multiple comparisons test. ns denotes not significant.

### 3.3 Hydroxytyrosol, a metabolote of oleuropein stimulates 5-HT secretion

When administered orally, oleuropein undergoes deglycosylation and hydrolysis during digestion, and is converted to hydroxytyrosol, which possesses high antioxidant activity (Figure 3A).^28^ Therefore, we investigated whether hydroxytyrosol stimulates 5-HT secretion. The results showed that hydroxytyrosol stimulated 5-HT secretion via Ca²⁺ influx without cytotoxicity (Figure 3B to D). Furthermore, the effects of hydroxytyrosol on 5-HT secretion and Ca²⁺ influx were abolished by the use of inhibitors of T-type and L-type voltage-dependent Ca^2+^ channels (Figure 3E to H). These results suggest that the ability of oleuropein to induce 5-HT secretion is primarily attributed to the tyrosol moiety, which remains intact even after digestion.

**Figure 3.**
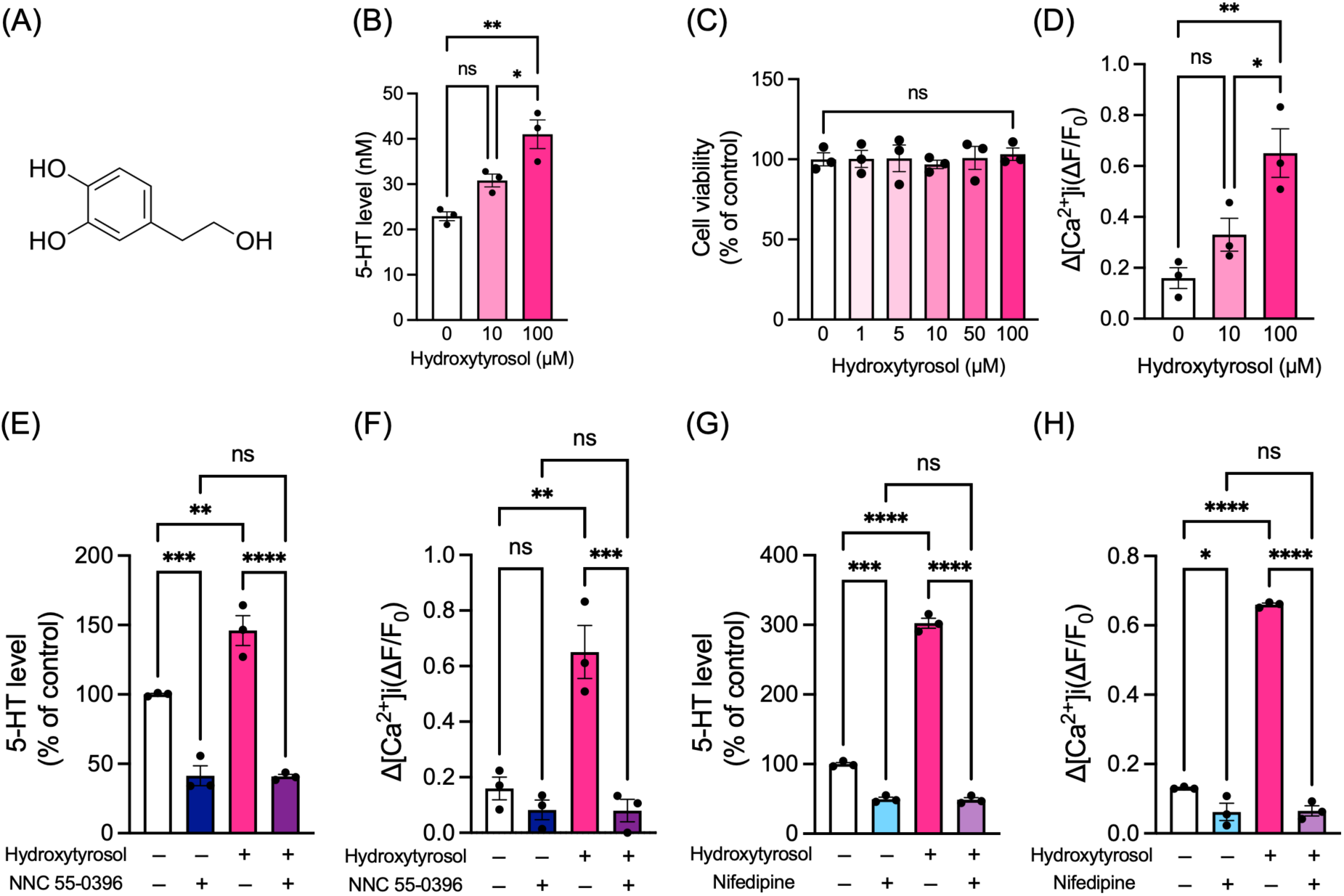
Effects of hydroxytyrosol on 5-HT secretion from enterochromaffin-like cells and Ca^2+^ influx. (**A**) Structure of hydroxytyrosol. (**B**) Dose-dependent effects of hydroxytyrosol on 5-HT secretion. QGP-1 cells were incubated with HBSS containing different concentrations of hydroxytyrosol, and the supernatant was subjected to HPLC analysis. (**C**) Dose-dependent effects of hydroxytyrosol on cell viability. QGP-1 cells were incubated with HBSS containing different concentrations of hydroxytyrosol for 60 min and subjected to the WST-8 assay. (**D**) Dose-dependent effects of hydroxytyrosol on cytosolic Ca^2+^ levels. Fluo-8 AM was internalized into QGP-1 cells, and hydroxytyrosol was added. The fluorescence intensity was then measured over time for 5 min. Ionomycin was used as a positive control. The average change in fluorescence intensity from 0 min after hydroxytyrosol addition to each time point was expressed as Δ[Ca^2+^]i (ΔF/F0). (**E**) Effects of T-type voltage-dependent Ca^2+^ channel inhibitor on 5-HT secretion by hydroxytyrosol. QGP-1 cells were incubated with HBSS containing 100 µM hydroxytyrosol for 5 min with or without NNC 55-0396. (**F**) Effects of T-type voltage-dependent Ca^2+^ channel inhibitor on Ca^2+^ influx induced by hydroxytyrosol. Fluo-8 AM was internalized into QGP-1 cells and hydroxytyrosol was added with or without NNC 55-0396. The fluorescence intensity was then measured over time for 5 min, and the average change in fluorescence intensity from 0 min after hydroxytyrosol addition to each time point was expressed as Δ[Ca^2+^]i (ΔF/F0). (**G**) Effects of L-type voltage-dependent Ca^2+^ channel inhibitor on 5-HT secretion by hydroxytyrosol. QGP-1 cells were incubated with HBSS containing 100 µM hydroxytyrosol for 5 min, with or without nifedipine. (**H**) Effects of L-type voltage-dependent Ca^2+^ channel inhibitor on Ca^2+^ influx induced by hydroxytyrosol. Fluo-8 AM was internalized into QGP-1 cells, and hydroxytyrosol was added with or without nifedipine. The fluorescence intensity was then measured over time for 5 min and the average change in fluorescence intensity from 0 min after hydroxytyrosol addition to each time point was expressed as Δ[Ca^2+^]i (ΔF/F0). Bars represent the mean ± SEM, and dots indicate individual data points (*n* = 3). \**p* < 0.05, \*\**p* < 0.01, \*\*\**p* < 0.001, \*\*\*\**p* < 0.0001, one-way ANOVA, Tukey’s multiple comparisons test. ns denotes not significant.

### 3.4 Oleuropein stimulates 5-HT secretion from mouse gut tissue

Next, we investigated whether oleuropein could stimulate 5-HT secretion from the gut. We first identified regions within the gut where enterochromaffin cells are abundant by utilizing immunofluorescence staining of Swiss-rolled mouse intestinal sections with antibodies specific to 5-HT and chromogranin A (CHGA), an enterochromaffin cell marker. The signals derived from 5-HT and CHGA were frequently observed in the proximal colon, indicating a relatively high number of enterochromaffin cells in this region (Figure 4A). Then, we isolated proximal colon tissue from mice and cultured it in the presence of oleuropein. The results showed that the addition of oleuropein significantly increased 5-HT secretion into the culture supernatant (Figure 4B). Furthermore, this 5-HT-secreting effect was significantly abolished by co-treatment with inhibitors of T-type and L-type voltage-dependent Ca²⁺ channels (Figure 4B). These results indicate that oleuropein stimulates 5-HT secretion via voltage-dependent Ca²⁺ channels in the mouse gut, as it does in QGP-1 cells.

**Figure 4.**
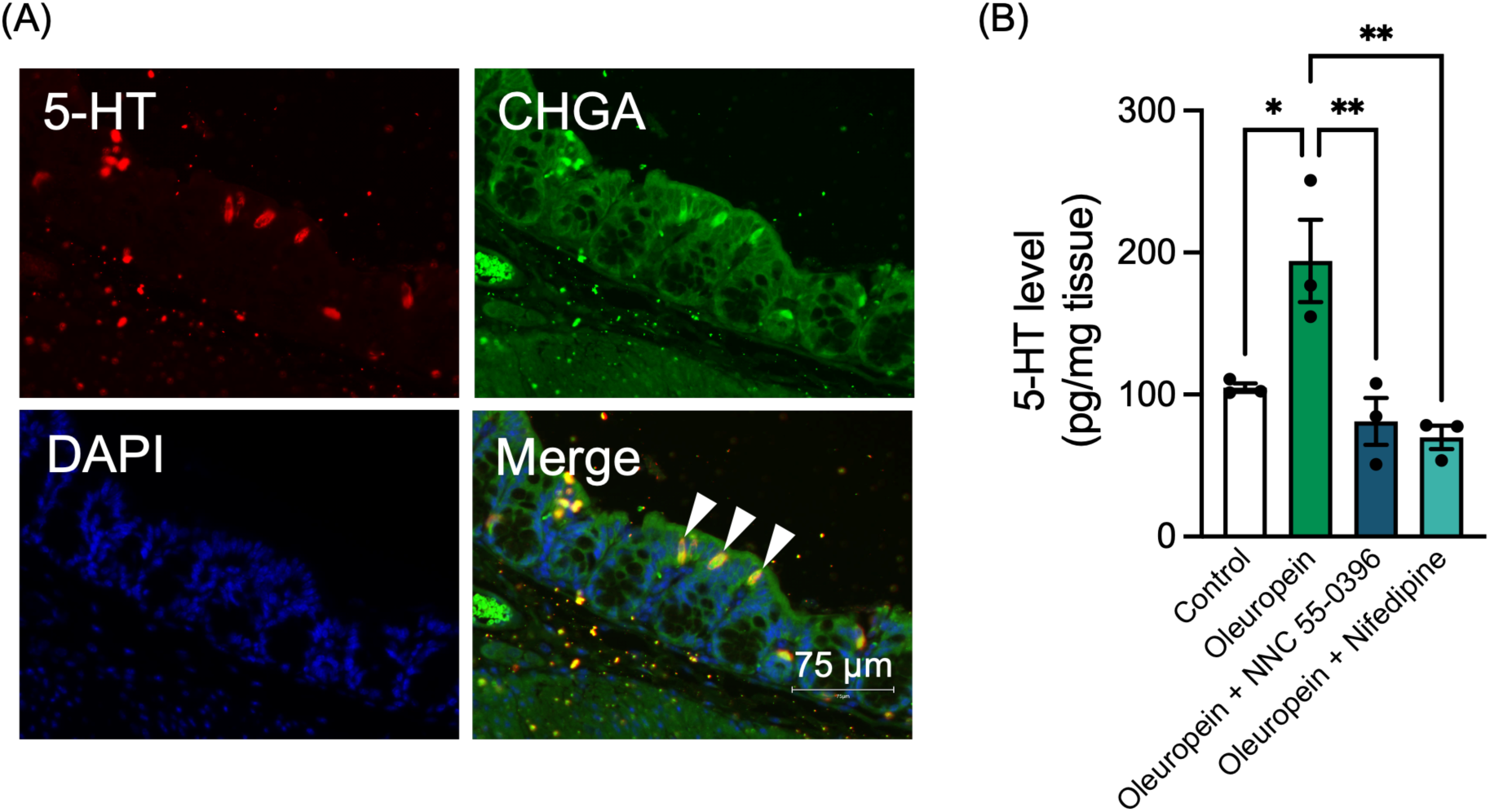
Effects of oleuropein on 5-HT secretion from mouse colon tissue. (**A**) Identification of enterochromaffin cells in the mouse digestive tract. Paraffin sections of Swiss-rolled mouse digestive tract were deparaffinized and incubated with antibodies for 5-HT and CHGA (enterochromaffin cell marker) after antigen-activated. After incubation with secondary antibodies conjugated with Cy2 or Cy3, the sections were covered with a mounting medium containing DAPI, and fluorescent images were acquired. The images show a magnified view of the colon region. White arrowheads indicate the enterochromaffin cells. (**B**) Effects of oleuropein on 5-HT secretion from mouse colon tissue. The proximal side of the colon tissue was dissected and washed with HBSS containing fluoxetine. Then, colon tissues were incubated for 10 min with HBSS containing fluoxetine with or without oleuropein, NNC 55-0396 and nifedipine. The culture supernatant was subjected to HPLC analysis. Bars represent the mean ± SEM, and dots indicate individual data points (*n* = 3). \**p* < 0.05, and \*\**p* < 0.01, one-way ANOVA, Tukey’s multiple comparisons test.

### 3.5 Oleuropein acutely regulates mental function in a 5-HT_3_ receptor-dependent manner

Recent studies have shown that enterochromaffin cells and peripheral 5-HT can regulate central functions.^7,8^ Based on our findings that oleuropein stimulates the secretion of 5-HT from the gut, we investigated the effects of oleuropein administration on mental functions. Mice were orally administered oleuropein once and underwent a series of behavioral tests to assess the acute effects on mental functions (Figure 5A). Furthermore, to confirm whether the effect of oleuropein on mental function was mediated by 5-HT, we used ramosetron, an inhibitor of the 5-HT_3_ receptor, which is deeply involved in the gut-brain axis.^29,30^ The results showed no significant changes due to oleuropein administration in the open field test, which examines emotionality and anxiety, and tail suspension test, which examines depression-like behavior (Figure 5B to D). On the other hand, in the forced swim test which examines depression-like behavior, oleuropein administration significantly increased the immobility time (Figure 5E). Importantly, oleuropein did not increase the immobility time when administered after administration of ramosetron. These results indicate that acute oral intake of oleuropein affects the central functions in a 5-HT_3_ receptor signal-dependent manner.

**Figure 5.**
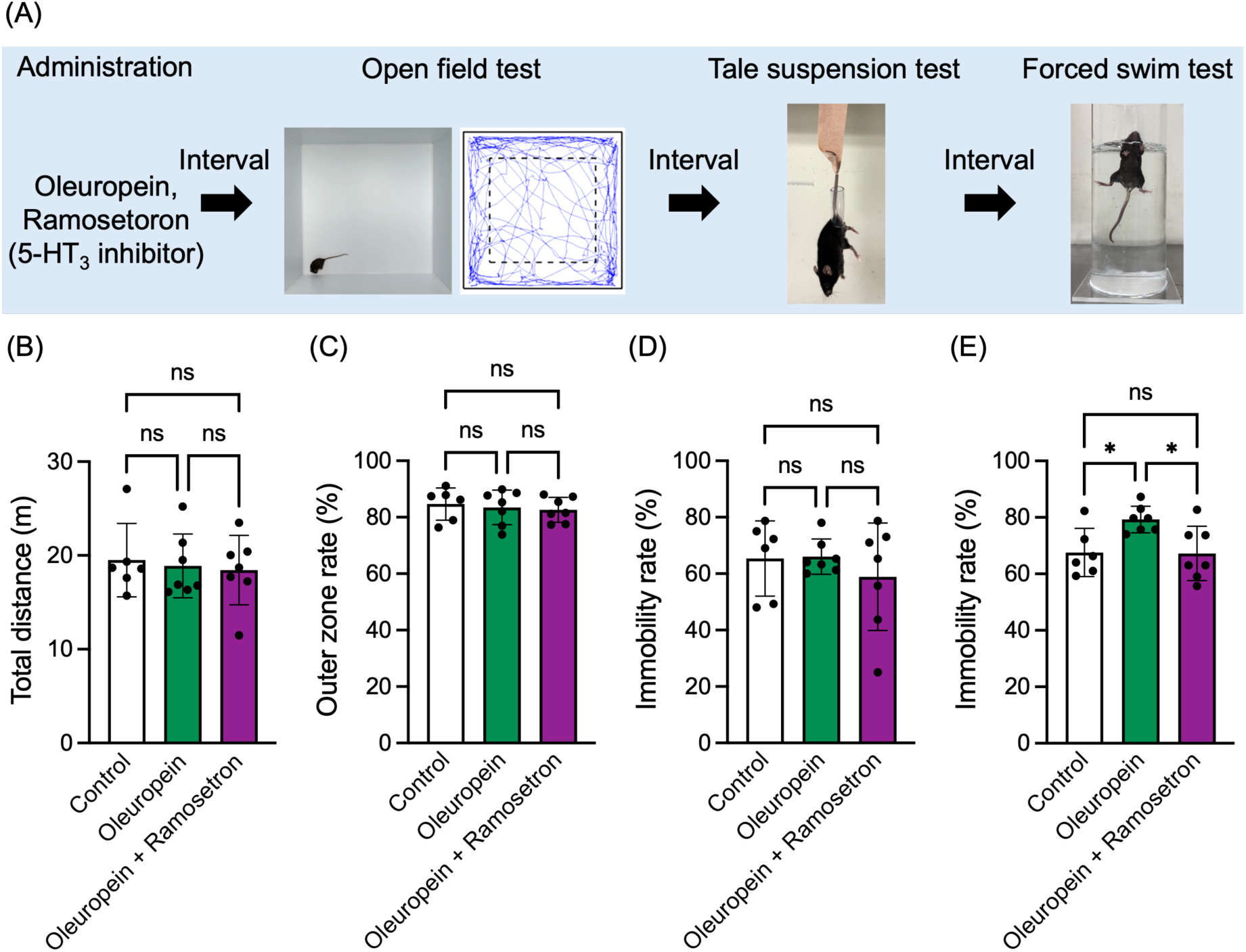
Acute effects of oral administration of oleuropein on mental function in mice. (**A**) Schematic diagram of the behavioral tests procedure. Mice were acclimated to the test room and administered oleuropein or ramosetron orally and then subjected to behavioral tests in the following order: open-field test, tail-suspension test, and forced swim test. (**B**) Effects of oral administration of oleuropein on the total distance traveled in the open field test. A single mouse was placed in the center of the arena, and its locomotor activity was recorded from above for 5 min. The videos were analyzed using the Mouse Activity software to calculate the total distance traveled. (**C**) Effects of oral administration of oleuropein on the time spent in the outer area of the arena in the open field test. A single mouse was placed in the center of the arena, and its locomotor activity was recorded from above for 5 min. The videos were analyzed using the Mouse Activity software to calculate the time spent in the outer area of the arena. (**D**) Effects of oral administration of oleuropein on the immobility time in the tale suspension test. Mouse was suspended by attaching tape to the tip of its tail. The activity was recorded for 6 min, and the immobility time during the 5 min period excluding the first 1 min was accumulated. (**E**) Effects of oral administration of oleuropein on the immobility time in the forced swim test. A single mouse was placed in an acrylic pipe with a water-filled base. The activity was recorded for 6 min, and the immobility time during the 5 min period excluding the first 1 min was accumulated. Bars represent the mean ± SEM, and dots indicate individual data points (*n* = 6-7). \**p* < 0.05, one-way ANOVA, Tukey’s multiple comparisons test. ns denotes not significant.

## 4. DISCUSSION

This study demonstrated that oleuropein stimulates 5-HT secretion from the gut and thereby regulates mental function (Figure 6). Since no food factors that regulate mental function via gut 5-HT secretion have been reported to our knowledge, our findings provide important evidence from the perspective of dietary regulation of mental function via the gut-brain axis. Furthermore, since olive consumption is known to be associated with the prevention and treatment of 5-HT-related conditions, such as mental and gastrointestinal disorders, our findings suggest that the 5-HT secretion induced by oleuropein may play a role in the physiological effects of olives.

**Figure 6.**
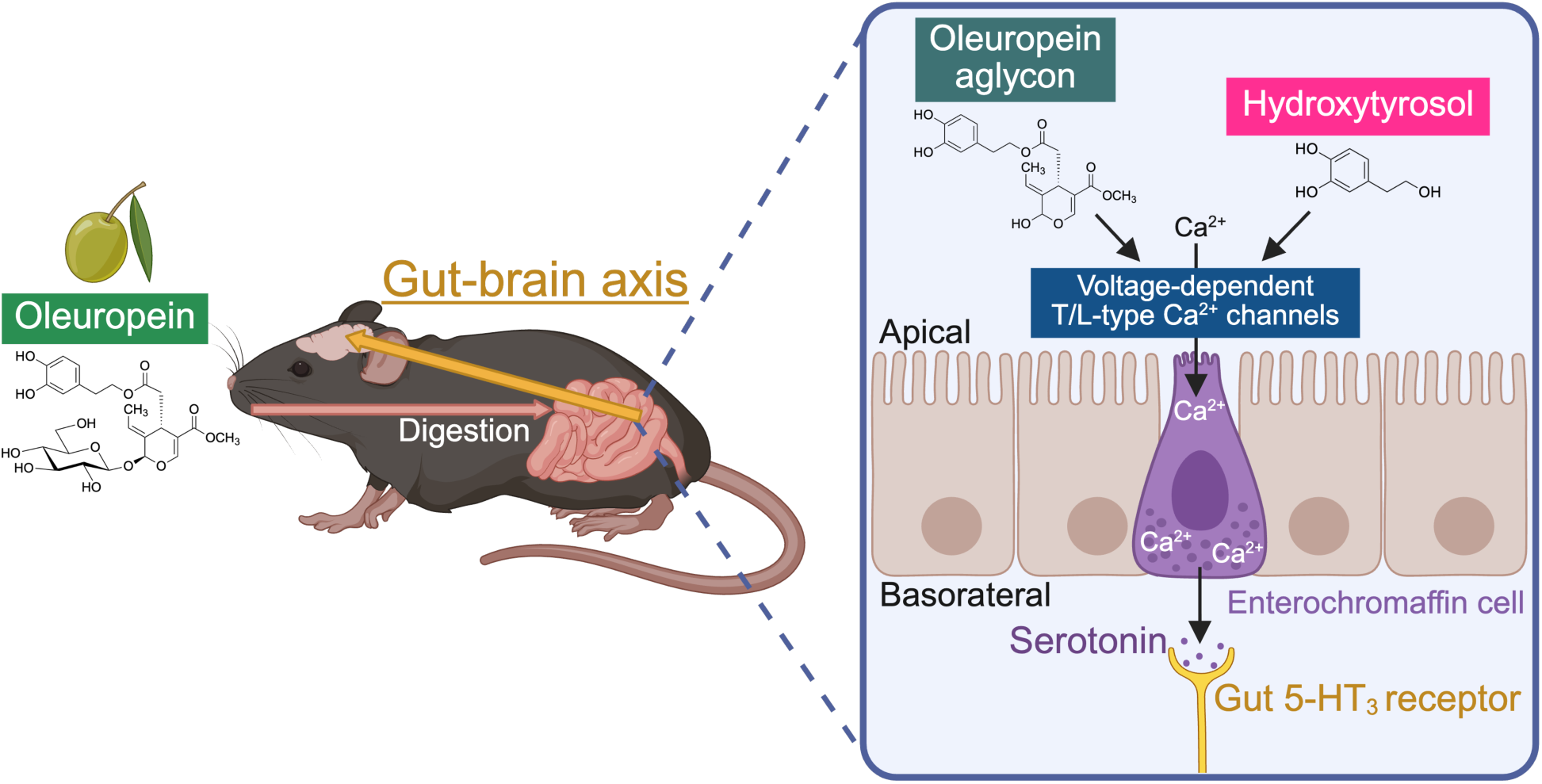
A schematic diagram illustrating the effects of oleuropein-induced 5-HT secretion from enterochromaffin cells on central function. After oral administration, oleuropein reaches the lower digestive tract as an aglycone or hydroxytyrosol metabolite. Enterochromaffin cells detect the tyrosol moiety of oleuropein aglycon or hydroxytyrosol, causing voltage-dependent Ca^2+^ channels to open and promoting Ca^2+^ influx into the cells. This stimulates 5-HT secretion, which acts on 5-HT_3_ receptors in the gut and influences mental function via the vagus nerve afferent. Created in BioRender. Kamei, Y. (2026) https://BioRender.com/gw4ai88

Our findings show that oleuropein stimulates 5-HT secretion from enterochromaffin-like cells and mouse colon through Ca^2+^ influx via not only T-type voltage-dependent Ca^2+^ channel but also L-type voltage-dependent Ca^2+^ channels. Considering that L-type voltage-dependent Ca^2+^ channels are expressed on the apical side of enterochromaffin cells,^26,27^ these results confirm the efficacy of orally ingested oleuropein on gut 5-HT secretion. Recently, oleuropein was identified as a potent activator of the mitochondrial Ca^2+^ uniporter in skeletal muscle among 5,000 bioactive molecules.^31^ Similarly, many polyphenols have been reported to improve the mitochondrial membrane potential.^32–34^ However, few studies have examined their direct effects on the membrane potential of cell membranes. Since mitochondrial Ca^2+^ uniporters are activated by local increases in Ca^2+^ levels,^35,36^ the cell membrane-mediated influx of Ca^2+^ that we have identified in this study may serve as the starting point for oleuropein’s effects on mitochondria. Consistent with this, mitochondria are located adjacent to Ca^2+^ release units containing L-type voltage-dependent Ca²⁺ channels in the T-tubules of skeletal muscle.^37^ Further research is needed to elucidate the effects of oleuropein-induced Ca^2+^ influx in intestinal epithelial cells on Ca^2+^ dynamics in mitochondria and the endoplasmic reticulum.

Voltage-dependent Ca²⁺ channels open in response to changes in membrane potential caused by ion flux across the plasma membrane.^38^ Therefore, it is highly likely that oleuropein first alters the membrane potential via some molecules on the cell membrane surface, resulting in the opening of voltage-dependent Ca^2+^ channels. For example, lipids have been shown to bind to Cd1d present in enterochromaffin cell membranes, causing potassium ion fluctuations that open the voltage-dependent Ca^2+^ channel, ultimately enhancing 5-HT secretion.^39^ Although TRPA1 was the leading candidate in the mechanism by which enterochromaffin cells detect food factors,^14^ this study revealed that oleuropein does not act via TRPA1. Contrary to our results, one study have shown that oleuropein acts on TRPA1 and TRPV1.^40^ The contradiction between these results may be due to the influence of oxides that are secondarily generated after polyphenols containing oleuropein are dissolved in water. A previous study showed that epigallocatechin gallate acted on TRPA1 and TRPV1 when left for several hours after preparation, rather than immediately after preparation.^41^ In this study, the solution was prepared immediately before use, and the effects were examined within a very short time after administration; therefore, the effects of oxidation products were not observed. Therefore, it is possible that secondary oxidation products which are generated from oleuropein act on TRP channels; however, experiments with longer time scales are required to confirm this. Further research using comprehensive analytical methods is needed to identify the initial target molecules acted upon by oleuropein before the opening of the voltage-dependent Ca^2+^ channels.

In this study, we demonstrated that acute oral administration of oleuropein enhances depression-like behavior in a 5-HT_3_ dependent manner. Since 5-HT_3_ receptors have been shown to be expressed in neurons adjacent to enterochromaffin cells,^10^ and are also therapeutic targets for irritable bowel syndrome, they may play a crucial role in the gut-brain axis via the vagus nerve afferent.^10,42^ Mice with specifically activated enterochromaffin cells have been reported to exhibit anxiety induced in a 5-HT_3_ signal-dependent manner.^8^ Together these findings indicate that gut 5-HT has the potential to regulate central functions, and that the control of gut 5-HT by food factors may play an important role in improving mental disorders. However, the present results contradict several previous reports that administration of oleuropein improves depression-like behavior.^43,44^ One possible reason for this discrepancy is the difference in administration routes. In most previous studies oleuropein was administered intraperitoneally, which may have made it more likely to act on the brain than in this study. Previous reports on curcumin, a type of polyphenol, suggest that intraperitoneal administration results in significantly higher blood concentrations and increased transfer to the brain than oral administration.^45^ Although oleuropein is unlikely to cross into the brain due to its structure, it is possible that hydroxytyrosol, a relatively small molecule produced by oleuropein, crosses into the brain and acts on T-type voltage-dependent Ca^2+^ channels expressed in the brain.^46,47^ Another possible reason for the discrepancy with previous reports on the effects of oleuropein on cognitive function is the difference in the duration of administration. Many prior studies employed chronic administration, potentially inducing numerous secondary effects. Notably, chronic oleuropein administration has been shown to affect the gut microbiota,^48,49^ making the impact on gut bacteria non-negligible. The results obtained in this study under the physiological conditions of oral administration and through acute testing suggest that the effects on the brain and the involvement of gut microbiota are considered negligible.

This study has several limitations. First, we did not analyze the direct acute effects of oleuropein intake on the brain. Oleuropein does not readily cross the blood-brain barrier, and given the experimental design of this acute study, it is unlikely that oleuropein directly affects the brain. However, to demonstrate the gut-to-brain mental control mechanisms mediated by food factors, it is necessary to investigate whether stimulation of the gut by oleuropein induces a brain response and to clarify the causal relationship between the gut and the brain. Second, while oral administration of a 5-HT_3_ receptor antagonist *in vivo* diminished the effects of oleuropein on central function, this drug may have affected the brain. Further studies using mice with gut-specific knockout of 5-HT-related factors, such as 5-HT_3_ receptors, or selective vagal deafferentation, are needed to confirm the effect. Despite these limitations, we identified oleuropein as a novel potent stimulant of gut 5-HT and modulator of central function via the 5-HT_3_ receptor. These findings provide a basis for future therapeutic strategies for mental disorders by food factors which target gut 5-HT.

## ABBREVIATIONS

AM: acetoxymethyl ester
BAPTA: 1,2-Bis(2-aminophenoxy)ethane-*N*,*N*,*N*’,*N*’-tetraacetic acid
CHGA: chromogranin A
DMSO: dimethyl sulfoxide
FBS: fetal bovine serum
HBSS: Hank’s balanced salt solution
HPLC: high performance liquid chromatography
TBS: tris buffer saline
TPH: tryptophan hydroxylase
TRP: transient receptor potential
WST: water soluble tetrazolium
5-HT: 5-hydroxytryptamine.

## Author contributions

Y.K. conceptualized the study, acquired funding, developed the methodology, conducted animal experiments, analyzed data, visualized data, supervised the project, and wrote the original draft. R.S. conducted cell and *ex vivo* experiments, formal analysis, and data curation. N.O. and M.A. conducted animal experiments, formal analysis, and data curation. T.F. developed the methodology, reviewed and edited the manuscript. M.A. conceptualized the study, developed the methodology, provided resources, supervised the project, reviewed and edited the manuscript.

## Notes

The authors declare no competing financial interest.

## ACKNOWLEDGEMENTS

The authors are grateful to Dr. S. Kaihara for linguistic advice. This study was supported by Support Center for Advanced Medical Sciences, Tokushima University Graduate School of Biomedical Sciences.

